# Sex Differences in Aortic Remodeling During Blood Pressure Reduction Following Gradual Hypertension Development in Spontaneously Hypertensive Rats

**DOI:** 10.64898/2026.08.17.745137

**Authors:** Yuki Hayashi, Yoshihiro Ujihara, Masanori Nakamura, Shukei Sugita

## Abstract

Cardiovascular disease risk is higher in men than in women. Although sex differences in aortic wall adaptation following antihypertensive treatment have been reported in acute hypertension models, the response after gradually developing hypertension, which mimics human essential hypertension, remains unclear. This study investigated sex differences in aortic wall adaptation following acute blood pressure reduction after gradually developing hypertension.

**Method:** Seventeen-week-old spontaneously hypertensive rats (SHRs) were assigned to the Hypertensive group or the antihypertensive (Reversal) group (*N* = 5/sex each). The Reversal group received the antihypertensive drug captopril for 4 weeks to maintain systolic blood pressure below 130 mmHg. Age-matched Wistar Kyoto rats (*N* = 3/sex) served as normotensive (Normal) group. After the experimental period, arterial wall thickness, circumferential wall stress, smooth muscle cell phenotype, and histological changes were evaluated.

**Results:** Antihypertensive treatment significantly reduced systolic blood pressure in both sexes. Both male and female SHRs exhibited elevated circumferential wall stress during the gradual development of hypertension. In females, antihypertensive treatment significantly reduced medial thickness compared with the Hypertensive group, whereas males showed no reduction. Circumferential wall stress in female Reversal group did not differ significantly from either the Hypertensive or Normal group, whereas males exhibited a significant reduction in circumferential wall stress compared with the Hypertensive group. Furthermore, the reduced collagen area fraction in the Hypertensive group returned to the normotensive levels only in females following antihypertensive treatment.

**Conclusion:** These findings indicate that vascular remodeling induced by gradually developing hypertension is more effectively reversed by antihypertensive treatment in females than in males.

## Introduction

Hypertension is a leading cause of premature mortality^1^ and a major risk factor for cardiovascular diseases, including aortic dissection^2^ and aortic aneurysm^3^. The prevalence of hypertension exceeds 60% among men aged ≥ 50 years and women aged ≥ 60 years in Japan.^4^ The primary goal of current antihypertensive therapy is to reduce cardiovascular risk through blood pressure lowering.^5^ However, normalization of blood pressure does not necessarily restore vascular function to a healthy state. Experimental studies have shown that pulse wave velocity, a marker of arterial stiffness, remains elevated even after blood pressure normalization.^6^ Furthermore, impairments in endothelium-dependent vasodilation and smooth muscle cell (SMC) function may persist despite normalized blood pressure.^6^ These observations suggest that normalization of blood pressure does not necessarily restore vascular homeostasis.

Biological tissues adapt structurally to changes in mechanical loading. According to Roux’s law, ^7^ increased mechanical loading induces hypertrophy, whereas reduced loading leads to tissue atrophy. This concept has also been applied to the arterial wall. Indeed, in experimental models of acute hypertension induced by renal artery clipping, the aortic wall thickens within approximately two weeks.^8^ Such thickening is considered a functional adaptation that reduces the elevated circumferential wall stress associated with increased blood pressure, thereby restoring mechanical homeostasis.

Sex differences have been reported in the reversibility of this adaptive remodeling. In a renal artery clipping model, female rats exhibited arterial wall thinning and restoration of circumferential wall stress toward normotensive levels after normalization of blood pressure, whereas these responses were absent in male rats. ^9^ These findings suggest sex-specific differences in the restoration of vascular mechanical homeostasis, although the underlying mechanisms remain unclear. Understanding such differences is clinically important because aortic dissection and aortic aneurysm occur more frequently in men than in women.^10–12^

Vascular remodeling in response to hypertension varies according to the mode and rate of blood pressure elevation. In the angiotensin II infusion model, a widely used model of acute hypertension, arterial wall thickening reduced circumferential wall stress toward normotensive levels.^13,14^ In contrast, a different pattern is observed in the spontaneously hypertensive rat (SHR), a model of gradually developing hypertension that shares many characteristics with human essential hypertension.^15–17^ Although arterial wall thickening also occurs in SHR, this adaptation is insufficient to restore circumferential wall stress to normotensive levels.^18^ These observations indicate that vascular adaptation to gradually developing hypertension differs fundamentally from that observed in acute hypertension. However, despite the clinical relevance of gradually developing hypertension, it remains unclear whether rapid pharmacological blood pressure reduction can re-establish arterial mechanical homeostasis and whether this response differs between males and females. Restoration of arterial mechanical homeostasis is expected to involve structural remodeling of the vessel wall, including alterations in extracellular matrix (ECM) composition and vascular SMC phenotype.

The aim of this study was to investigate aortic wall adaptation to rapid blood pressure reduction following the gradual development of hypertension. We assessed the restoration of circumferential wall stress and associated structural remodeling, including alterations in ECM composition and vascular SMC phenotype. We also examined whether these adaptive responses differ between males and females.

## 2. Materials and Methods

### 2.1 Experimental Animals and Blood Pressure Models

Fifteen-week-old male and female SHRs were used in this study. Age-matched male and female Wistar Kyoto (WKY) rats served as normotensive controls. All animal procedures were approved by the Institutional Animal Care and Use Committee of our institute (approval numbers: D-2025006 and D-2025007).

WKY rats served as the Normal group (*N* = 3/sex), whereas SHRs were assigned to either the Hypertensive group (*N* = 5/sex) or the Reversal group (*N* = 5/sex). In the Reversal group, captopril, an angiotensin-converting enzyme inhibitor, was administered in drinking water beginning at 17 weeks of age. The dose (50–300 mg/kg/day) was adjusted to maintain systolic blood pressure (SBP) below approximately 130 mmHg.

The time required to achieve the target SBP (<130 mmHg) ranged from approximately 1 to 4 weeks. Animals were then maintained under this condition for an additional 4 weeks before experimentation. Consequently, age at euthanasia ranged from 21 to 25 weeks. This 4-week interval was defined as the blood pressure reduction period. Animals in the Hypertension group were euthanized at ages matched to those of the corresponding animals in the Reversal group.

Female SHR were given free access to a high-salt diet containing 4% NaCl (Oriental Yeast, Tokyo, Japan) to promote the development of hypertension, because female SHR exhibit lower blood pressure than male SHR.^15^ The high-salt diet was administered from 15 weeks of age until the end of the experiment in the Hypertension group and from 15 to 17 weeks of age, before the initiation of antihypertensive treatment, in the Reversal group. The overall experimental protocols for female and male rats are summarized in Supplemental Figure S1A– B.

### 2.2 Blood Pressure Measurement

SBP was measured at least weekly using the tail-cuff method with a noninvasive automated blood pressure monitoring system (BP-98A-L, Softron, Tokyo, Japan).

### 2.3 Preparation of Aortic Samples

The thoracic aorta was harvested as previously described.^19–21^ Briefly, rats were euthanized by gradual-fill CO_2_ inhalation, and death was confirmed before tissue collection. The thoracic aorta was excised in Krebs–Henseleit solution. While maintaining the *in vivo* axial length, markers were placed at 5-mm intervals along the vessel using gentian violet. A 20-mm segment of the descending thoracic aorta immediately above the diaphragm was used for analyses.

The excised aorta was pressure-fixed in 10% neutral-buffered formalin for 2 h using a custom-built pressurization system shown in Supplemental Figure S1C. The intraluminal fixation pressure was set to the mean SBP of each animal during the final 4 weeks before euthanasia.

### 2.4 Morphology and Mechanical Stress Analysis

After pressure fixation, transverse sections perpendicular to the vessel axis with a thickness of 150 μm were prepared and imaged using confocal laser scanning microscopy to evaluate medial elastic lamellae. Detailed imaging procedures are described in the Supplemental File.

All image analyses were performed using Fiji ImageJ software (version 1.54f; National Institutes of Health, Bethesda, MD, USA). Wall thickness was defined as the radial distance between the internal and external elastic laminae. Measurements were performed at three locations per image in four images per cross-section, yielding a total of 12 measurements per animal. The mean value was used as the wall thickness (*t*).

The luminal boundary was manually traced from whole-cross-sectional images of the aorta, and the luminal area (*S*) at the mean SBP (*P*) of each animal was measured. Assuming a circular lumen, the luminal radius at SBP (*R*) was calculated as:

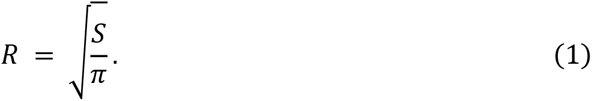

The radius-to-thickness ratio *R*/*t* was calculated as an index of morphological adaptation of the vessel wall to altered blood pressure.

Circumferential wall stress at SBP (*σ*_θSBP_) was calculated according to Laplace’s law:

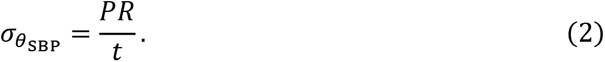

### 2.5 Smooth Muscle Cell Phenotype Evaluation

To assess the vascular SMC phenotype, transverse aortic sections were immunostained for the contractile SMC markers calponin and α-smooth muscle actin (α-SMA). Negative-control sections were processed without primary antibodies. Images of the medial layer were acquired by confocal laser scanning microscopy. Detailed methods are provided in the Supplemental File.

For each specimen, *z*-stack images were converted into two-dimensional images using maximum-intensity projection. Medial regions were identified from elastin autofluorescence, and the mean fluorescence intensity (MFI) of each marker (*I*_dye_) was quantified. To correct for variations in imaging conditions, negative-control sections from the same animal were imaged on the same day with identical acquisition settings, and the corresponding medial MFI was defined as *I*_NC_. Normalized fluorescence intensity (*I*_NORM_) was calculated as:

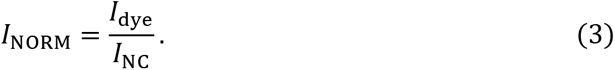

### 2.6 Extracellular Matrix Composition Evaluation

Pressure-fixed thoracic aortic specimens were stained with Elastica van Gieson (EVG) for elastin, Azan for collagen, and hematoxylin and eosin (H&E) for cellular nuclei and tissue morphology. Histological sections were imaged by light microscopy for quantitative analysis. Detailed imaging procedures are described in the Supplemental File.

Histological components of the medial layer were quantified using a color-based classification method.^22,23^ A custom ImageJ macro classified EVG-stained images into elastin, collagen, cytoplasm, and background, and Azan-stained images into collagen, muscle fibers, and background according to representative RGB values for each component. After excluding background pixels, the area fraction of each component within the medial layer was quantified. The area fractions of elastin and collagen were used for subsequent analyses.

To assess changes in medial SMC number, SMCs within the medial layer were quantified in H&E-stained sections.

### 2.7 Statistical Analysis

Data are presented as mean ± standard deviation (SD). Statistical analyses were performed using two-way analysis of variance (ANOVA) with sex (male and female) and blood pressure condition (Normal, Hypertension, and Reversal) as factors. When the main effect of blood pressure condition was significant, post hoc comparisons were performed using the Tukey–Kramer test. Analyses were conducted in R (version 4.6.0; R Foundation for Statistical Computing, Vienna, Austria). A two-sided *p* < 0.05 was considered statistically significant.

## 3. Results

### 3.1 Blood Pressure Profiles

Mean SBP during antihypertensive treatment period is shown in Figure 1. Two-way ANOVA revealed significant main effects of blood pressure condition (*p* < 0.001) and sex (*p* = 0.016) on mean SBP, with no significant interaction (*p* = 0.698). Post hoc analyses showed that SBP was significantly higher in the Hypertension group than in the Normal and Reversal groups in both sexes. These results confirm successful induction of hypertension and normalization of SBP following antihypertensive treatment.

**Figure 1.**
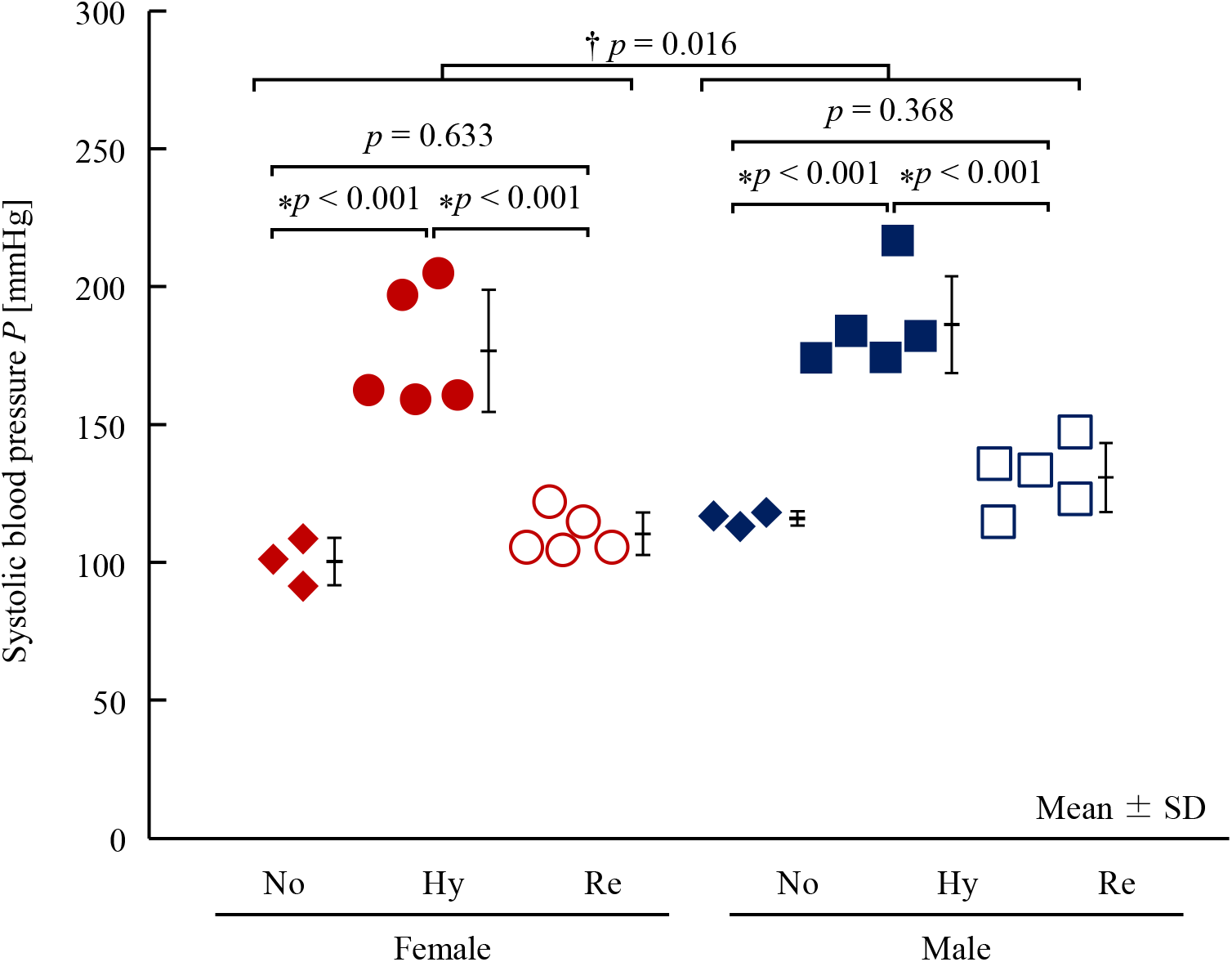
Systolic blood pressure in each group. Each dot represents an individual animal. ⁎: *p* < 0.05 by Tukey–Kramer test. †: *p* < 0.05 for the main effect of sex by two-way ANOVA. No, Normal; Hy, Hypertension; Re, Reversal.

### 3.2 Wall Thickness and Lumen Radius-to-Wall Thickness Ratio

Representative images of pressure-fixed aortas at the mean SBP during the antihypertensive treatment period are shown in Figure 2A–F. In females, aortic wall thickness appeared greater in the Hypertension group (Figure 2B) and lower in the Reversal group (Figure 2C). In males, wall thickness was similar between the Normal and Hypertension groups (Figure 2D, E), whereas lower values were observed in the Reversal group (Figure 2F).

**Figure 2.**
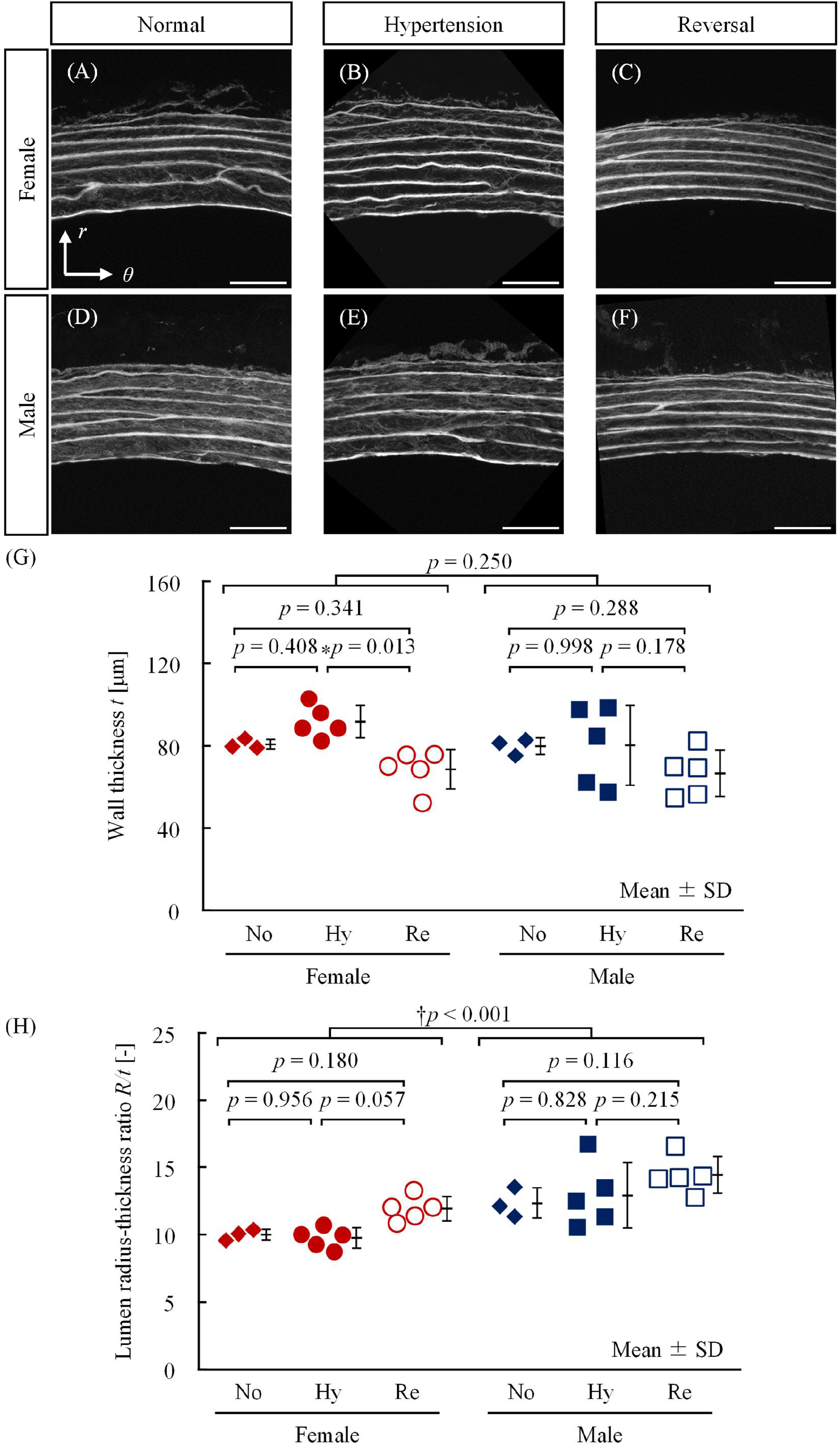
Morphological parameters of the aortic wall in female and male rats. (A–F) Representative cross-sectional images of the aortic wall from (A–C) female and (D–F) male rats under (A, D) Normal, (B, E) Hypertension, and (C,F) Reversal conditions. Scale bars, 50 μm. *r*, radial; *θ*, circumferential directions. (G) Wall thickness (*t*). (H) Radius-to-thickness ratio (*R/t*). Each dot represents an individual animal. ⁎: *p* < 0.05 by Tukey–Kramer test. †: *p* < 0.05 for the main effect of sex by two-way ANOVA. No, Normal; Hy, Hypertension; Re, Reversal.

Quantitative measurements of aortic wall thickness are shown in Figure 2G. Two-way ANOVA revealed a significant main effect of blood pressure condition (*p* = 0.007), whereas neither the main effect of sex (*p* = 0.250) nor the interaction (*p* = 0.571) was significant. Post hoc analysis identified a significant difference only between the female Hypertension and Reversal groups, with lower wall thickness in the Reversal group. No significant differences were detected among the male groups.

The lumen radius-to-wall thickness ratio (*R*/*t*) is shown in Figure 2H. Two-way ANOVA revealed significant main effects of blood pressure condition (*p* = 0.010) and sex (*p* < 0.001), with no significant interaction (*p* = 0.798). However, post hoc analysis did not detect significant differences among blood pressure condition groups. In both sexes, *R*/*t* values were comparable between the Normal and Hypertension groups. Males exhibited significantly higher *R*/*t* values than females overall, independent of blood pressure condition.

### 3.3 Circumferential Stress

Circumferential wall stress at the mean SBP of each animal during the antihypertensive treatment period is shown in Figure 3. Two-way ANOVA revealed significant main effects of blood pressure condition (*p* < 0.001) and sex (*p* < 0.001), with no significant interaction (*p* = 0.693). In both females and males, circumferential wall stress was significantly higher in the Hypertension group than in the Normal group. Circumferential wall stress was significantly lower in the Reversal group than in the Hypertension group in males, whereas no significant difference was observed between these groups in females. No significant differences were detected between the Normal and Reversal groups in either sex. These results indicate that gradually developing hypertension increased circumferential wall stress in both sexes. Following blood pressure reduction, circumferential wall stress tended to decrease in females, although this reduction did not reach statistical significance, whereas males showed a significant decrease.

**Figure 3.**
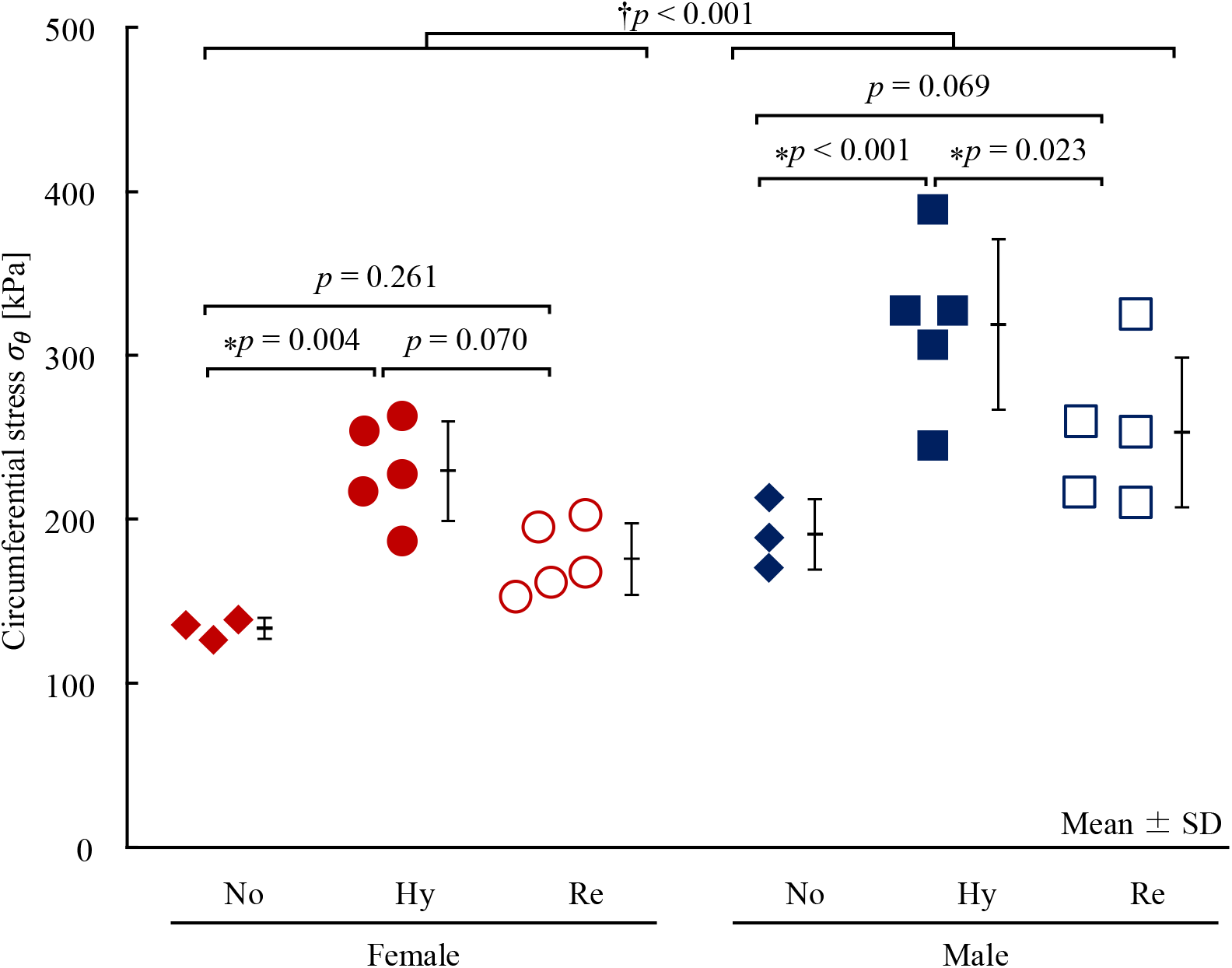
Circumferential wall stress at systolic blood pressure in each group. Each dot represents an individual animal. ⁎: *p* < 0.05 by Tukey–Kramer test. †: *p* < 0.05 for the main effect of sex by two-way ANOVA. No, Normal; Hy, Hypertension; Re, Reversal.

### 3.4 Normalized calponin fluorescence intensity

Representative immunofluorescence images of calponin are shown in Figure 4A–F. Quantification of normalized calponin MFI is shown in Figure 4G. Two-way ANOVA revealed a significant main effect of blood pressure condition (*p* = 0.040), whereas neither the main effect of sex (*p* = 0.330) nor the interaction was significant (*p* = 0.772). Post hoc analysis detected no significant differences among groups within either sex, although calponin MFI tended to be lower in the Hypertension and Reversal groups than in the Normal group in both sexes. In contrast, normalized α-SMA MFI was unaffected by blood pressure condition, sex, or their interaction (Supplemental Figure S2). These results suggest that calponin immunofluorescence intensity may be affected by blood pressure condition, whereas no significant effects on α-SMA immunofluorescence intensity were detected.

**Figure 4.**
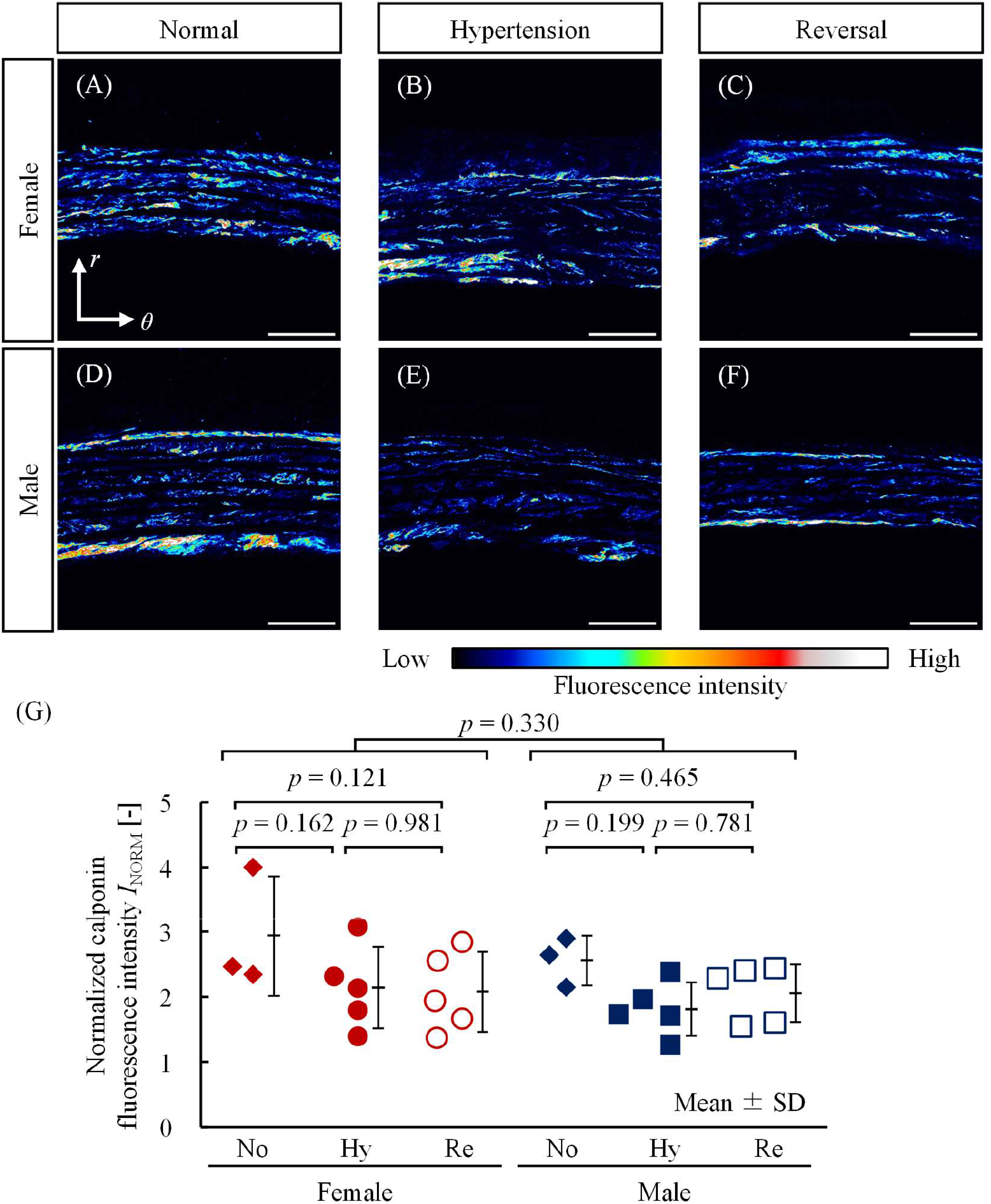
Immunofluorescent staining of calponin in each group. (A–F) Representative immunofluorescence images of the aortic wall from (A–C) female and (D–F) male rats under (A, D) Normal, (B, E) Hypertension, and (C, F) Reversal conditions. Scale bars, 50 μm. *r*, radial; *θ*, circumferential directions. The luminal side appears at the bottom of each image. (G) Normalized calponin fluorescence intensity (*I*_NORM_). Each dot represents an individual animal. ⁎: *p* < 0.05 by Tukey–Kramer test. †: *p* < 0.05 for the main effect of sex by two-way ANOVA. No, Normal; Hy, Hypertension; Re, Reversal.

### 3.5 Elastin Area fraction

Representative EVG-stained cross-sections of the aorta are shown in Figure 5A–F. The elastic fiber area fraction quantified is shown in Figure 5G. Two-way ANOVA revealed a significant main effect of blood pressure condition (*p* < 0.001), whereas neither the main effect of sex (*p* = 0.238) nor the interaction was significant (*p* = 0.285). In females, the elastic fiber area fraction was significantly lower in the Hypertension group than in the Normal group, with no significant differences involving the Reversal group. In males, both the Hypertension and Reversal groups exhibited significantly lower elastic fiber area fractions than the Normal group, whereas no significant difference was observed between the Hypertension and Reversal groups. Elastic fiber area fraction decreased in response to hypertension. The difference between the Normal and Reversal groups was no longer significant in females, whereas reduced values persisted in males.

**Figure 5.**
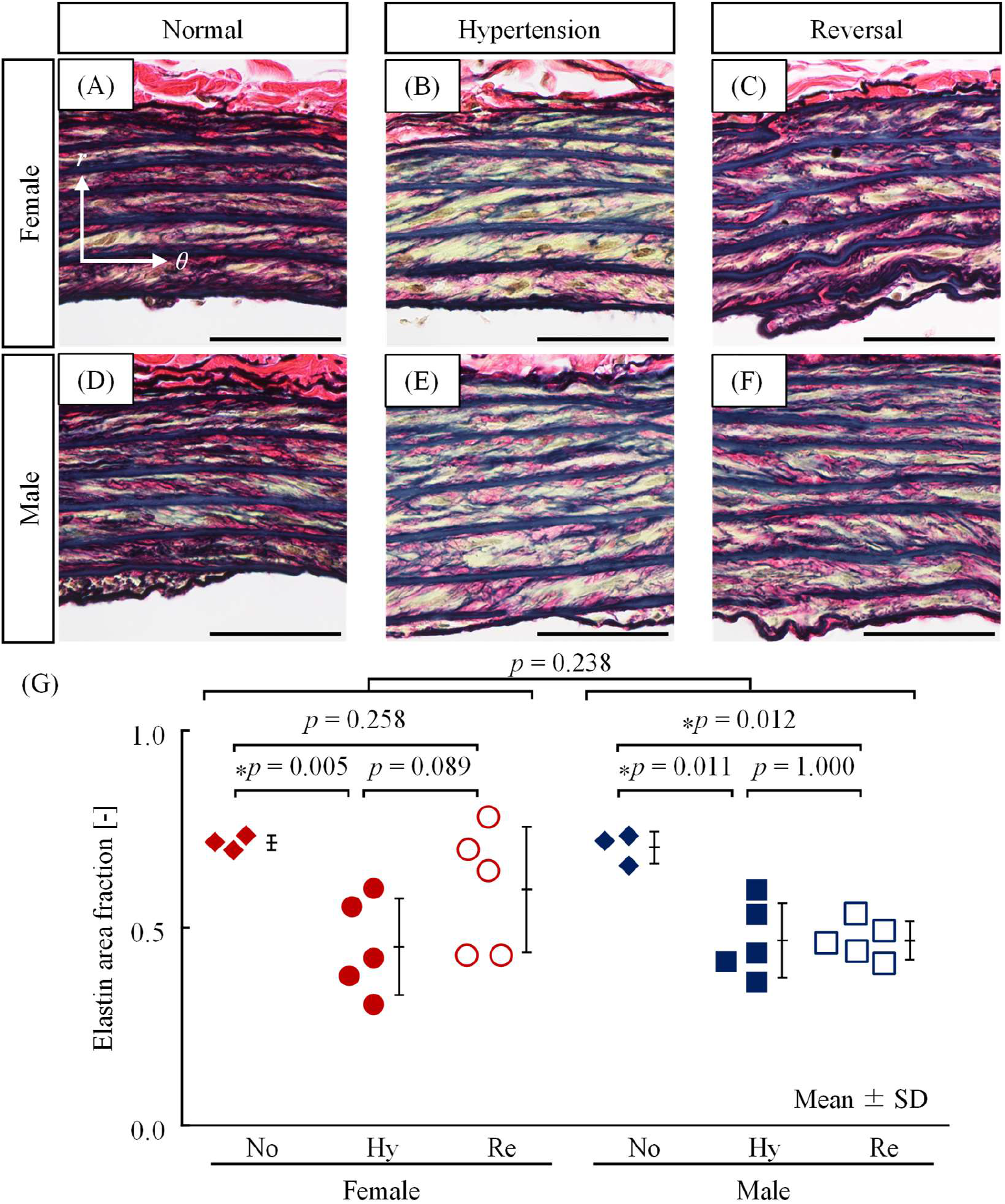
Elastica Van Gieson (EVG) staining of the aortic wall in each group. Elastic fibers are stained dark purple. (A–F) Representative EVG-stained cross-sectional images of the aortic wall from (A–C) female and (D–F) male rats under (A, D) Normal, (B, E) Hypertension, and (C, F) Reversal conditions. Elastic fibers are stained dark purple. Scale bars, 50 μm. *r*, radial; *θ*, circumferential directions. (G) Elastin area fraction. Each dot represents an individual animal. ⁎: *p* < 0.05 by Tukey–Kramer test. No, Normal; Hy, Hypertension; Re, Reversal.

### 3.6 Collagen Area fraction

Representative Azan-stained cross-sections of the aorta are shown in Figure 6A–F. The medial collagen fiber area fraction is shown in Figure 6G. Two-way ANOVA revealed significant main effects of blood pressure condition (*p* = 0.001) and sex (*p* = 0.006), whereas no significant interaction was observed (*p* = 0.583). In females, the collagen fiber area fraction was significantly lower in the Hypertension group than in both the Normal and Reversal groups. In males, the Hypertension group exhibited a significantly lower collagen fiber area fraction than the Normal group, with no significant difference between the Hypertension and Reversal groups. Collagen fiber area fraction decreased in response to hypertension in both sexes and returned to levels comparable to the Normal group following blood pressure reduction, with a more pronounced response in females. Overall, males exhibited lower collagen fiber area fractions than females.

**Figure 6.**
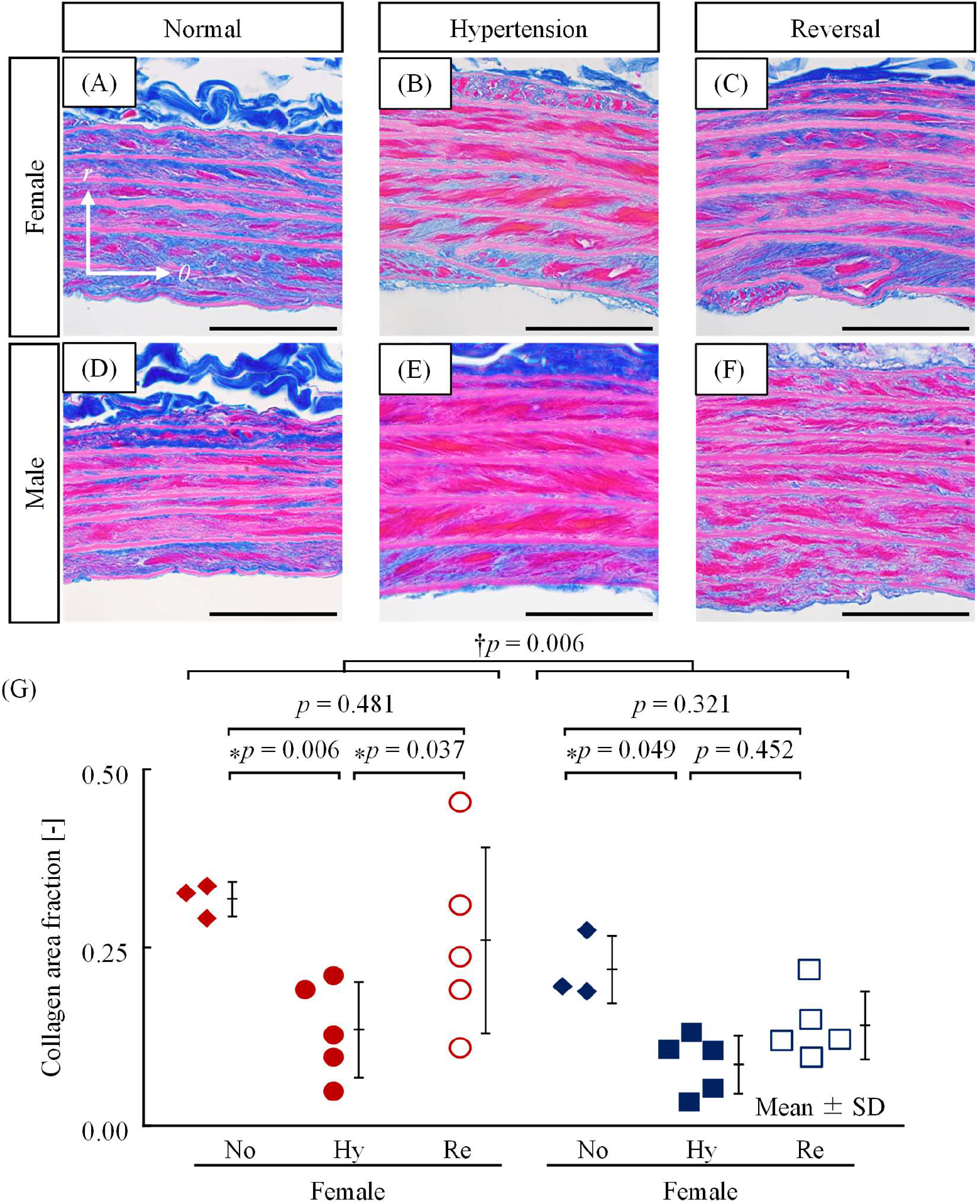
Azan staining of the aortic wall in each group. Collagen fibers are stained blue. (A– F) Representative Azan-stained cross-sectional images of the aortic wall from (A–C) female and (D–F) male rats under (A, D) Normal, (B, E) Hypertension, and (C, F) Reversal conditions. Collagen fibers are stained blue. Scale bars, 50 μm. *r*, radial; *θ*, circumferential directions. (G) Collagen area fraction. Each dot represents an individual animal. ⁎: *p* < 0.05 by Tukey–Kramer test. †: *p* < 0.05 for the main effect of sex by two-way ANOVA. No, Normal; Hy, Hypertension; Re, Reversal.

### 3.7 Number of cells

Representative H&E-stained cross-sections of the aorta are shown in Supplemental Figure S3A–F. The number of medial SMCs is shown in Supplemental Figure S3G. Two-way ANOVA revealed no significant main effects of blood pressure condition (*p* = 0.420) or sex (*p* = 0.735), and no significant interaction (*p* = 0.769), indicating no detectable differences in medial SMC number among groups.

## 4. Discussion

In the present study, we evaluated aortic wall adaptation to hypertension and subsequent blood pressure reduction in SHR, a model of gradually developing hypertension.^15,16^ The major findings were as follows. First, both male and female SHR in the hypertension group of the present study exhibited persistently elevated circumferential wall stress, unlike the normalization of wall stress reported in acute hypertension models, indicating incomplete restoration of mechanical homeostasis. Second, following short-term antihypertensive treatment, wall thickness decreased significantly only in females, whereas males showed no significant wall thinning but exhibited a significant decrease in circumferential wall stress. Third, aortic elastin and collagen content, reduced by hypertension in both sexes, tended to recover after treatment only in females. Together, these findings suggest that vascular adaptation to gradually developing hypertension differs from that in acute hypertension models and that the response to blood pressure normalization is influenced by sex.

Previous studies have shown that vascular remodeling responses differ depending on the mode of blood pressure elevation. In models of acute hypertension, arterial wall thickening restores circumferential wall stress to normotensive levels.^8^ In both female and male SHR in this study, wall thickness tended to increase in the hypertension group (Figure 2), but this increase was insufficient to normalize circumferential wall stress (Figure 3), in line with previous reports in male SHR.^18^ Consistent with this interpretation, the *R*/*t* ratio remained comparable between the normotensive and hypertensive groups in both sexes (Figure 2H), despite the increase in blood pressure. These findings indicate limited morphological adaptation to hypertension in SHR, regardless of sex.

Circumferential wall stress was not normalized, yet histological remodeling was evident. Both male and female hypertensive rats exposed to gradual blood pressure elevation exhibited reduced elastin and collagen area fractions compared with normotensive rats (Figures 5 and 6). A previous study reported decreased elastin content in both sexes after acute hypertension,^9^ suggesting that elastin reduction is independent of the mode of blood pressure elevation. In contrast, collagen content was unchanged in both sexes in an acute hypertension model^9^, suggesting that gradual blood pressure elevation reduces collagen content. These ECM changes occurred despite persistent elevation of circumferential wall stress, indicating that vascular remodeling can proceed independently of the classical stress-homeostasis response.

Following blood pressure reduction after gradually developing hypertension, only females exhibited wall thinning together with relative preservation of circumferential wall stress (Figures 2G and 3). Despite the absence of a significant difference in the *R*/*t* ratio between the Hypertension and Reversal groups, the female Reversal group showed a tendency toward a higher *R*/*t* ratio (Figure 2H) associated with wall thinning (Figure 2G), suggesting structural adaptation that attenuated the stress reduction. No such tendency was observed in males. These observations indicate that structural adaptation following blood pressure reduction occurs more readily in females. Wolinsky reported that, following rapid blood pressure reduction after acute hypertension, females exhibited wall thinning accompanied by maintenance of circumferential wall stress, whereas males did not.^9^ Together, these findings suggest that females may have a greater propensity for stress homeostasis following rapid blood pressure reduction, regardless of whether hypertension develops gradually or abruptly.

Under hypertensive conditions, calponin, a contractile marker of vascular SMCs, tended to decrease in both sexes (Fig. 4). Previous studies have shown that cyclic equibiaxial stretch (1 Hz, 16%) simulating the mechanical environment of human hypertension induces phenotypic switching of cultured SMCs from a contractile to a synthetic phenotype.^24^ In addition, vascular SMCs derived from patients with hypertension have been reported to exhibit a synthetic phenotype.^25^ These findings support the present results. In contrast, α-SMA expression was unchanged (Supplemental Figure S2), suggesting that SMCs did not complete a transition to the synthetic phenotype but rather exhibited partial phenotypic modulation. Furthermore, calponin expression was not fully restored in either sex after 4 weeks of antihypertensive treatment (Figure 4), suggesting persistence of a partially modulated phenotype despite blood pressure normalization. No apparent sex differences were observed in the expression of either calponin or α-SMA.

Elastin and collagen area fractions tended to recover after blood pressure reduction only in females (Figures 5 and 6), although they decreased in both sexes under hypertensive conditions. These findings suggest greater ECM remodeling capacity in females than in males. Estrogen is known to regulate both ECM degradation and synthesis. Estrogen deficiency increases the expression of elastin- and collagen-degrading MMPs in the aorta,^26,27^ whereas estrogen promotes elastin synthesis in SMCs.^28^ Therefore, recovery of the elastin area fraction observed only in females (Figure 5) may reflect enhanced elastin synthesis together with reduced elastin degradation mediated by estrogen. The collagen area fraction also tended to recover only in females after blood pressure reduction (Figure 6); however, estrogen has not been consistently shown to promote collagen synthesis, suggesting that additional mechanisms may contribute to this recovery.

Circumferential wall stress at SBP was consistently higher in males than in females under all blood pressure conditions (Figure 3). This finding indicates that circumferential wall stress is not maintained at the same level between sexes in the gradually developing hypertension model, SHR. Similarly, Wolinsky reported higher circumferential wall stress in males than in females after 20 weeks of hypertension induced by renal artery clipping in Carworth rats, an acute hypertension model.^8^ Thus, despite differences in the mode of blood pressure elevation, both studies demonstrated higher circumferential wall stress in males. Furthermore, both the present study and Wolinsky’s study demonstrated higher circumferential wall stress in males under normotensive conditions.^9^ Taken together, these findings suggest that the higher circumferential wall stress observed in males is not unique to SHR and may reflect sex-specific differences in the target level of circumferential wall stress.

The *R*/*t* ratio of the rat thoracic aorta has been shown to increase progressively from 4 to 52 weeks of age, accompanied by a corresponding rise in circumferential wall stress.^29^ Body growth may be viewed as a process of gradual and sustained elevation of circumferential wall stress, analogous to hypertension development in SHR. Interestingly, Wolinsky reported that circumferential wall stress was not normalized during growth under normotensive conditions,^9^ and the present study showed a similar lack of normalization during gradually developing hypertension in SHR. Collectively, these observations raise the possibility that arterial adaptation is influenced not only by the absolute magnitude of circumferential wall stress but also by the rate at which it changes. A rapid increase in circumferential wall stress may activate stress-homeostatic mechanisms involving wall thickening, whereas a gradual increase, such as that associated with growth or hypertension in SHR, may allow a higher level of circumferential wall stress to become established as a new homeostatic set point. Taken together, these findings suggest that the homeostatic level of circumferential wall stress is not a universal constant but rather a dynamic set point influenced by factors such as sex, growth, and the rate of blood pressure change.

## Perspectives

This study demonstrated sex differences in the mechanical adaptation and histological remodeling of the aortic wall during antihypertensive treatment following gradually developing hypertension, a condition that more closely resembles human essential hypertension. The lack of circumferential wall stress normalization during gradually developing hypertension previously observed in males was also evident in females, contrasting with the normalization of circumferential wall stress reported in acute hypertension in both sexes. However, antihypertensive treatment normalized circumferential wall stress in females but not in males, indicating sex-specific differences in vascular adaptation to antihypertensive treatment. In addition, recovery of elastin and collagen area fractions was less pronounced in males than in females. Together, these findings suggest that vascular adaptive responses are influenced by both the mode of blood pressure elevation and biological sex. Future studies should identify the mechanical and molecular mechanisms governing these responses. Since these sex differences were observed under hemodynamic conditions resembling human essential hypertension, the findings may contribute to the development of sex-specific strategies for antihypertensive therapy and vascular protection.

## Novelty and Relevance

### What is new?

- Circumferential wall stress remained elevated during gradually developing hypertension in male and female SHRs.
- Four weeks of antihypertensive treatment induced medial thinning and stress normalization only in females.
- Recovery of collagen and elastin area fractions was greater in females.
- Circumferential wall stress was consistently higher in males.

### What is relevant?

- Gradually developing hypertension induces limited morphologic but consistent histological remodeling.
- Vascular recovery after antihypertensive treatment differs by sex.
- Blood pressure trajectories may redefine the circumferential wall stress set point.

### Clinical/pathophysiological

- Early blood pressure control and sex-specific treatment strategies may improve vascular recovery in chronic hypertension.

## Nonstandard Abbreviations and Acronyms

ANOVA: analysis of variance
ECM: extracellular matrix
H&E: hematoxylin and eosin
MFI: mean fluorescence intensity
SBP: systolic blood pressure
SD: standard deviation
SHR: spontaneously hypertensive rat
SMC: smooth muscle cell
WKY: Wistar Kyoto

## References

1. Wang C, Yuan Y, Zheng M, Pan A, Wang M, Zhao M, Li Y, Yao S, Chen S, Wu S, et al. Association of age of onset of hypertension with cardiovascular diseases and mortality. J Am Coll Cardiol. 2020;75:2921–2930.

2. Nienaber CA, Fattori R, Mehta RH, Richartz BM, Evangelista A, Petzsch M, Cooper JV, Januzzi JL, Ince H, Sechtem U, et al. Gender-related differences in acute aortic dissection. Circulation. 2004;109:3014–3021.

3. Ailawadi G, Eliason JL, Upchurch GR Jr. Current concepts in the pathogenesis of abdominal aortic aneurysm. J Vasc Surg. 2003;38:584–588.

4. Hisamatsu T, Segawa H, Kadota A, Ohkubo T, Arima H, Miura K. Epidemiology of hypertension in Japan: beyond the new 2019 Japanese guidelines. Hypertens Res. 2020;43:1344–1351.

5. Umemura S, Arima H, Arima S, Asayama K, Dohi Y, Hirooka Y, Horio T, Hoshide S, Ikeda S, Ishimitsu T, et al. The Japanese Society of Hypertension Guidelines for the Management of Hypertension (JSH 2019). Hypertens Res. 2019;42:1235–1481.

6. Steppan J, Jandu S, Savage W, Wang H, Kang S, Narayanan R, Nyhan D, Santhanam L. Restoring blood pressure in hypertensive mice fails to fully reverse vascular stiffness. Front Physiol. 2020;11:824.

7. Kivell TL. A review of trabecular bone functional adaptation: what have we learned from trabecular analyses in extant hominoids and what can we apply to fossils?. J Anat. 2016;228:569–594.

8. Matsumoto T, Hayashi K. Stress and strain distribution in hypertensive and normotensive rat aorta considering residual strain. J Biomech Eng. 1996;118:62–73.

9. Wolinsky H. Effects of hypertension and its reversal on the thoracic aorta of male and female rats. Morphological and chemical studies. Circ Res. 1971;28:622–637.

10. Chang CY, Wu CF, Muo CH, Chang SS, Chen PC. Sex differences in temporal trends and risk factors of aortic dissection in Taiwan. J Am Heart Assoc. 2023;12:e027833.

11. Smedberg C, Steuer J, Leander K, Hultgren R. Sex differences and temporal trends in aortic dissection: a population-based study of incidence, treatment strategies, and outcome in Swedish patients during 15 years. Eur Heart J. 2020;41:2430–2438.

12. Pham MHC, Sigvardsen PE, Fuchs A, Kühl JT, Sillesen H, Afzal S, Nordestgaard BG, Køber LV, Kofoed KF. Aortic aneurysms in a general population cohort: prevalence and risk factors in men and women. Eur Heart J Cardiovasc Imaging. 2024;25:1235–1243.

13. Bersi MR, Bellini C, Wu J, Montaniel KRC, Harrison DG, Humphrey JD. Excessive adventitial remodeling leads to early aortic maladaptation in angiotensin-induced hypertension. Hypertension. 2016;67:890–896.

14. Humphrey JD. Mechanisms of vascular remodeling in hypertension. Am J Hypertens. 2021;34:432–441.

15. Fukuda S, Tsuchikura S, Iida H. Age-related changes in blood pressure, hematological values, concentrations of serum biochemical constituents and weights of organs in the SHR/Izm, SHRSP/Izm and WKY/Izm. Exp Anim. 2004;53:67–72.

16. Okamoto K, Tabei R, Fukushima M, Nosaka S, Yamori Y, Ichijima K, Haebara H, Matsumoto M, Maruyama T, Suzuki Y, et al. Further observations of the development of a strain of Spontaneously Hypertensive Rats. Jpn Circ J. 1966;30:703–716.

17. Elmarakby AA, Sullivan JC. Sex differences in hypertension: lessons from spontaneously hypertensive rats (SHR). Clin Sci (Lond). 2021;135:1791–1804.

18. Bézie Y, Lamazière JM, Laurent S, Challande P, Cunha RS, Bonnet J, Lacolley P. Fibronectin expression and aortic wall elastic modulus in spontaneously hypertensive rats. Arterioscler Thromb Vasc Biol. 1998;18:1027–1034.

19. Sugita S, Mizuno N, Ujihara Y, Nakamura M. Stress fibers of the aortic smooth muscle cells in tissues do not align with the principal strain direction during intraluminal pressurization. Biomech Model Mechanobiol. 2021;20:1003–1011.

20. Sugita S, Kato M, Wataru F, Nakamura M. Three-dimensional analysis of the thoracic aorta microscopic deformation during intraluminal pressurization. Biomech Model Mechanobiol. 2020;19:147–157.

21. Fukui W, Ujihara Y, Nakamura M, Sugita S. Direct visualization of interstitial flow distribution in aortic walls. Sci Rep. 2022;12:5381.

22. Kurihara G, Ujihara Y, Nakamura M, Sugita S. Delamination strength and elastin interlaminar fibers decrease with the development of aortic dissection in model rats. Bioengineering (Basel). 2023;10:1292.

23. Kurihara G, Ujihara Y, Nakamura M, Sugita S. Effect of elastin on delamination strength in the porcine aortic tunica media. J Biorheol. 2023;37:120–129.

24. Hu B, Song JT, Qu HY, Bi CL, Huang XZ, Liu XX, Zhang M. Mechanical stretch suppresses microRNA-145 expression by activating extracellular signal-regulated kinase 1/2 and upregulating angiotensin-converting enzyme to alter vascular smooth muscle cell phenotype. PLoS One. 2014;9:e96338.

25. Régent A, Ly KH, Lofek S, Clary G, Tamby M, Tamas N, Federici C, Broussard C, Chafey P, Liaudet-Coopman E, et al. Proteomic analysis of vascular smooth muscle cells in physiological condition and in pulmonary arterial hypertension: toward contractile versus synthetic phenotypes. Proteomics. 2016;16:2637–2649.

26. Horie K, Nanashima N, Maeda H, Tomisawa T, Oey I. Blackcurrant (Ribes nigrum L.) extract exerts potential vasculoprotective effects in ovariectomized rats, including prevention of elastin degradation and pathological vascular remodeling. Nutrients. 2021;13:560.

27. Miyamoto C, Kugo H, Hashimoto K, Moriyama T, Zaima N. Ovariectomy increases the incidence and diameter of abdominal aortic aneurysm in a hypoperfusion-induced abdominal aortic aneurysm animal model. Sci Rep. 2019;9:18330.

28. Natoli AK, Medley TL, Ahimastos AA, Drew BG, Thearle DJ, Dilley RJ, Kingwell BA. Sex steroids modulate human aortic smooth muscle cell matrix protein deposition and matrix metalloproteinase expression. Hypertension. 2005;46:1129– 1134.

29. Rachev A, Greenwald SE, Kane TP, Moore JE Jr, Meister JJ. Analysis of the strain and stress distribution in the wall of the developing and mature rat aorta. Biorheology. 1995;32:473–485.

