## Supplemental Methods and Figures for "Sex Differences in Aortic Remodeling During Blood Pressure Reduction Following Gradual Hypertension Development in Spontaneously Hypertensive Rats"

**Title**

**Shor title:** (no more than 50 characters including spaces)

Sex Differences in Aortic Remodeling Recovery

**Authors:**

Yuki Hayashi, B. Eng. †

Yoshihiro Ujihara, Ph. D. †

Masanori Nakamura, Ph. D. †, ††, §

Shukei Sugita, Ph. D. †, ††

**Affiliations:**

† Department of Electrical and Mechanical Engineering, Nagoya Institute of Technology, Gokiso-cho, Showa-ku, Nagoya 466-8555, Japan.

†† Center of Biomedical Physics and Information Technology, Nagoya Institute of Technology, Gokiso-cho, Showa-ku, Nagoya 466-8555, Japan.

§ Department of Nanopharmaceutical Sciences, Nagoya Institute of Technology, Gokiso-cho, Showa-ku, Nagoya 466-8555, Japan.

**Corresponding author:**

**Name:** Shukei Sugita

**Address:** Department of Electrical and Mechanical Engineering, Nagoya Institute of Technology, Gokiso-cho, Showa-ku, Nagoya 466-8555, Japan.

#### Expanded Methods

##### 1. Blood Pressure Measurement

SBP was measured using the tail-cuff method with a noninvasive automated blood pressure monitoring system (BP-98A-L, Softron, Tokyo, Japan). Measurements were performed at least once weekly between 15:00 and 17:00. Before each measurement session, rats were warmed at 38°C for 10 min. SBP was measured five consecutive times, and the highest and lowest values were excluded. The average of the remaining three measurements was used for analysis.

##### 2. Morphology and Mechanical Stress Analysis

The thoracic aorta was embedded in 3% agar and sectioned perpendicular to the vessel axis at a thickness of 150 μm using a linear slicer (NLS-MT, Dosaka-EM, Kyoto, Japan). Medial elastic lamellae were imaged using a confocal laser scanning microscope (FV3000, Olympus, Tokyo, Japan) equipped with a 60× oil-immersion objective lens (UPLSAPO60XO, Olympus). For each cross-section, four regions distributed circumferentially at approximately equal intervals were selected. Autofluorescence of the elastic lamellae was imaged using an excitation wavelength of 488 nm and an emission wavelength range of 500–540 nm. Whole-vessel cross-sectional images were also acquired using the same confocal laser scanning microscope equipped with a 10× objective lens (UPLSAPO10X2, Olympus).

**3. Immunofluorescence Staining and Confocal Microscopy for Smooth Muscle Cell Phenotype Evaluation**

To evaluate the vascular SMC phenotype, immunofluorescence staining was performed on aortic sections. Transverse sections (150 µm thick) prepared as described above were permeabilized with 0.2% Triton X-100 (MP Biomedicals, Santa Ana, CA, USA) in PBS(−) for 3 h. After permeabilization, the sections were blocked with 4% bovine serum albumin (BSA; A1470-25G, Sigma-Aldrich, Burlington, MA, USA) in PBS(−) for 1 h. Calponin (Rabbit Anti-calponin IgG Antibody, ab46794, Abcam, Cambridge, UK) and α-smooth muscle actin (α-SMA; Mouse Anti-α Smooth Muscle Actin Antibody, A5228, Sigma-Aldrich, Burlington, MA, USA) were used as primary antibodies. The sections were incubated with primary antibodies diluted 1:1000 in PBS(−) containing 0.2% BSA for 8 h at 37°C with shaking at 1 Hz. For secondary staining, Alexa Fluor 546 goat anti-rabbit IgG (A-11010, Thermo Fisher Scientific, Waltham, MA, USA) was used for calponin staining, and Alexa Fluor 546 goat anti-mouse IgG (A-11003, Thermo Fisher Scientific) was used for α-SMA staining. The sections were incubated with secondary antibodies diluted 1:1000 in PBS(−) containing 0.2% BSA under the same conditions as the primary antibodies and protected from light. Sections were washed three times with PBS(−) after each incubation step. Negative-control sections were processed without primary antibodies.

The medial layer of the stained aortic sections was imaged using a confocal laser scanning microscope (FV3000, Olympus) equipped with a 60× objective lens. Elastin autofluorescence (excitation: 488 nm; emission: 500–540 nm) and target protein fluorescence (excitation: 561 nm; emission: 570–670 nm) were acquired within the same field of view. Optical sections were obtained at intervals of 0.5–2 μm along the *z*-axis.

**4. Extracellular Matrix Composition Evaluation**

Tissue sectioning and histological staining were performed by Sapporo General Pathology Laboratory (Sapporo, Japan). Histological sections were imaged using an inverted microscope (IX73, Olympus) equipped with either a 40× objective lens (LUCPLFLN40X, Olympus) or a 60× objective lens (UPLSAPO60XW, Olympus). For each cross-section, images were acquired at four circumferential locations approximately 90° apart. Exposure time and white balance were determined for each specimen based on the first acquired image and were kept constant for all subsequent images of the same specimen.

First, representative RGB values were determined for each tissue component. In EVG-stained images, 30 pixels corresponding to elastin, collagen, cytoplasm, and background were manually selected, and the mean RGB value of each component was calculated as its representative RGB value. In Azan-stained images, representative RGB values were similarly determined for collagen, muscle fibers, and background. Next, for each pixel in the histological image, the sum of squared differences between the pixel RGB value and the representative RGB value of each component was calculated. Each pixel was then assigned to the component with the smallest sum of squared differences. Based on the resulting classification, the area fraction of each component was calculated as the ratio of the number of pixels assigned to that component to the total number of pixels within the medial layer, excluding background pixels. The area fractions of elastin and collagen were used for subsequent analyses.

### Supplemental Figures and Figure Legends

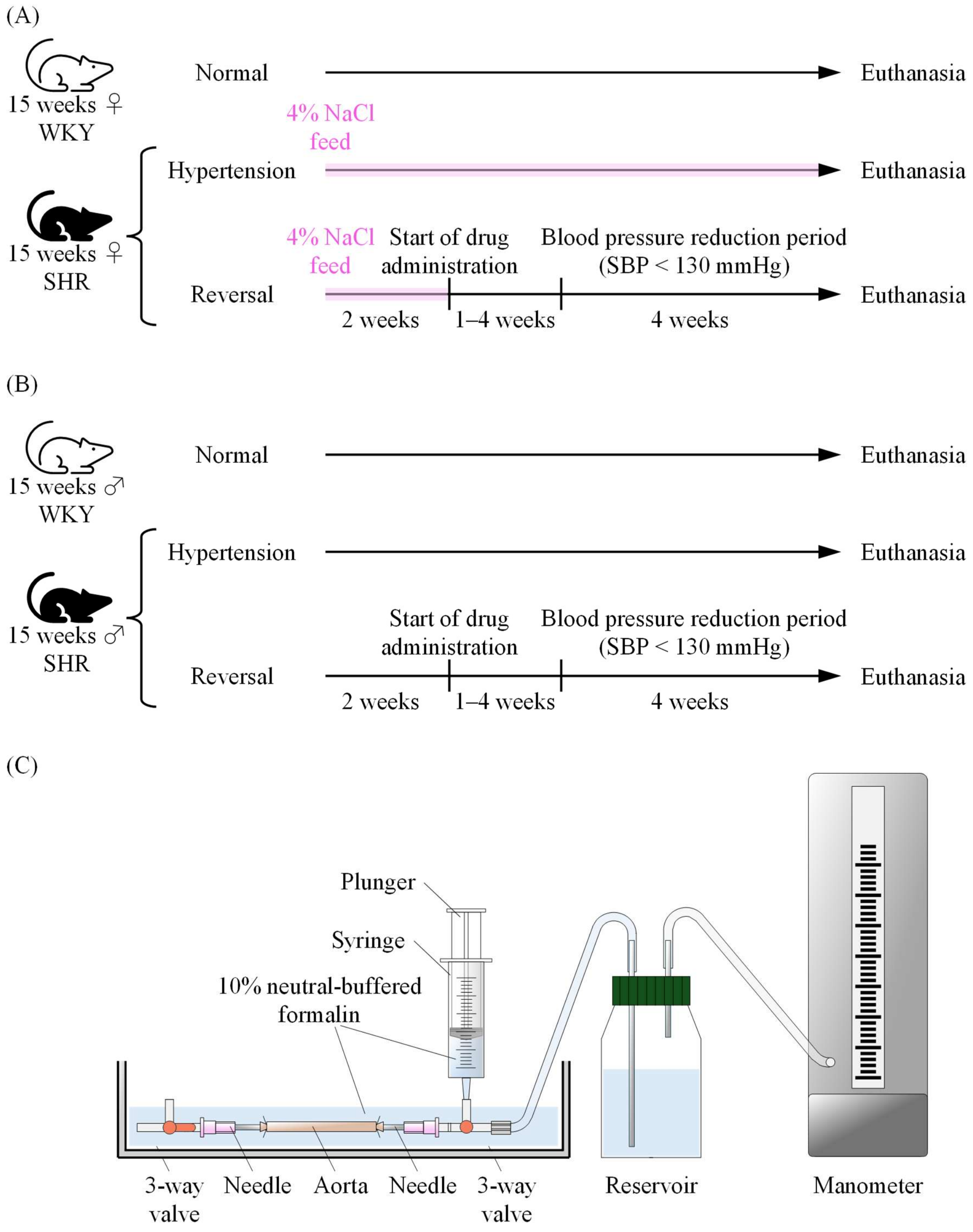


**Figure S1**. Experimental protocols and the custom-built pressurization system. (A) Protocol for female SHR and WKY rats. Female SHR were given free access to a high-salt diet containing 4% NaCl, as shown by the pink shading; in the reversal group, the diet was switched to a normal diet from 17 weeks of age. (B) Protocol for male SHR and WKY rats. In both (A) and (B), the reversal group received an antihypertensive drug in the drinking water from 17 weeks of age, and the period during which SBP was maintained at < 130 mmHg was defined as the blood pressure reduction period. (C) Custom-built pressurization system used for arterial fixation. A syringe was used to apply pressure equivalent to each animal's mean SBP during the blood pressure reduction period.

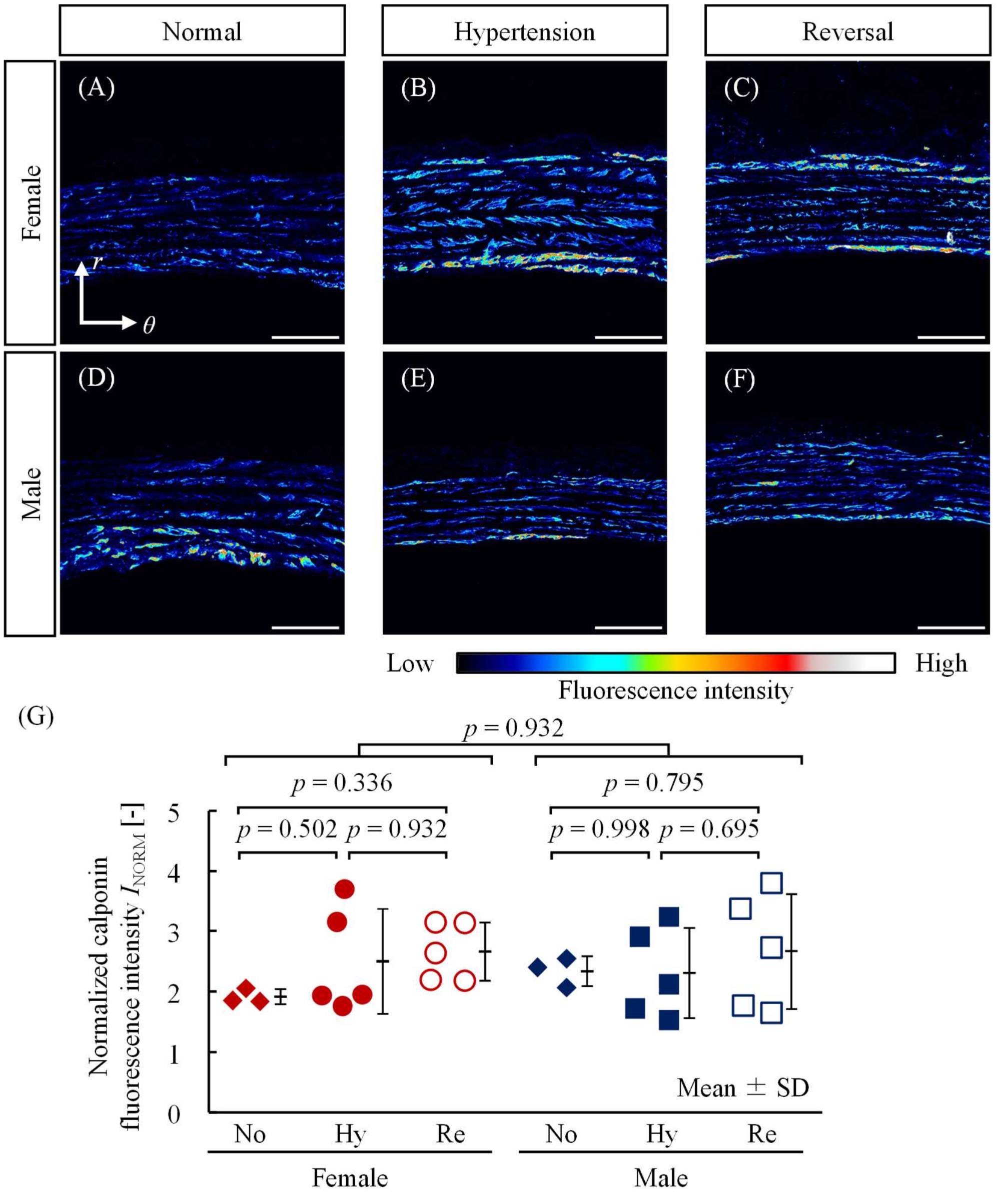


**Figure S2**. Immunofluorescent staining of α-SMA in each group. (A–F) Representative immunofluorescence images of the aortic wall from (A–C) female and (D–F) male rats under (A, D) Normal, (B, E) Hypertension, and (C, F) Reversal conditions. Scale bars, 50 μm. *r*, radial; *θ*, circumferential directions. The luminal side is located at the bottom of each image. (G) Normalized α-SMA fluorescence intensity ($I_{NORM}$). Each dot represents an individual animal. ∗: $p < 0.05$ by Tukey–Kramer test. †: $p < 0.05$ for the main effect of sex by two-way ANOVA. No, Normal; Hy, Hypertension; Re, Reversal.

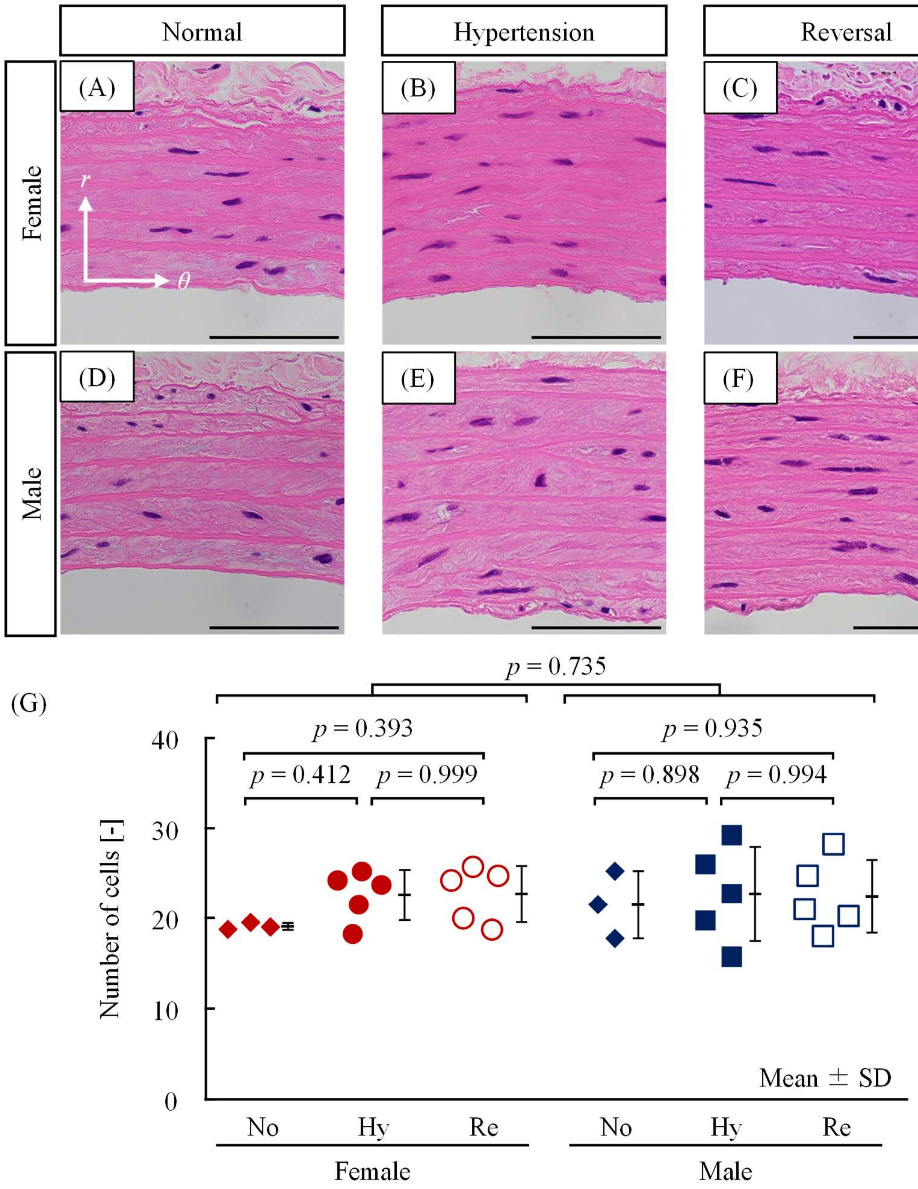


**Figure S3.** H&E staining of the aortic wall in each group. Cell nuclei are stained blue-purple. (A–F) Representative H&E-stained cross-sectional images of the aortic wall from (A–C) female and (D–F) male rats under (A, D) Normal, (B, E) Hypertension, and (C, F) Reversal conditions. Scale bars, 50 μm. *r*, radial; *θ*, circumferential directions. (G) Number of cells within the medial layer. Each dot represents an individual animal. ∗: $p < 0.05$ by Tukey–Kramer test. †: $p < 0.05$ for the main effect of sex by two-way ANOVA. No, Normal; Hy, Hypertension;

Re, Reversal.
